# Genetically-programmed Hypervesiculation of *Lactiplantibacillus plantarum* Increases Production of Bacterial Extracellular Vesicles with Therapeutic Efficacy in a Preclinical Inflammatory Bowel Disease Model

**DOI:** 10.1101/2025.06.20.660770

**Authors:** Nicholas H Pirolli, Daniel Levy, Alyssa Schledwitz, Natalia Sampaio Moura, Talia J. Solomon, Emily H. Powsner, Raith Nowak, Zuzanna Mamczarz, Christopher J. Bridgeman, Sulayman Khan, Laura Reus, Nidhi Anne, William E. Bentley, Jean-Pierre Raufman, Steven M. Jay

**Author notes:** Corresponding Author: Dr. Steven M. Jay, PhD. Conflict of interest: NHP, WEB and SMJ have intellectual property interests (pending US patent application) related to extracellular vesicle technology.

## Abstract

Inflammatory bowel diseases (IBD) affect over 6 million people globally and current treatments achieve only 10-20% rates of durable disease remission. Bacterial extracellular vesicles (BEVs) from probiotic lactic acid bacteria (LAB) are a promising novel therapeutic with mechanisms holding potential to drive increased rates of durable disease remission, including immunomodulation and intestinal epithelial tissue repair. However, translation of these cell-secreted nanovesicles is limited by long standing biomanufacturing hurdles, especially low production yields due to low biogenesis rates from cells. Here, our goal was to identify a candidate probiotic LAB that produces BEVs effective in a preclinical mouse model of IBD, and then genetically engineer the LAB for at least 10-fold increased production yields of BEVs, thereby passing a critical production threshold. We identified *Lactiplantibacillus plantarum* as a candidate LAB producing BEVs effective in treating acute dextran sulfate sodium (DSS)-induced murine colitis, and with greater efficacy than BEVs from probiotic *Escherichia coli* Nissle 1917. We then genetically engineered a hypervesiculating *L. plantarum* strain by inducible expression of a peptidoglycan-modifying enzyme, resulting in a 66-fold increase in BEV productivity. Finally, we confirmed hypervesiculating *L. plantarum* BEVs were therapeutically effective in the acute DSS mouse model of colitis and found these BEVs were superior in reducing mucosal tissue damage compared to live *L. plantarum* cells. These findings demonstrate that BEVs from genetically engineered hypervesiculating strain of *L. plantarum* are a promising preclinical therapeutic candidate for IBD that overcomes historical biomanufacturing limitations of BEV therapeutics.

## Introduction

Inflammatory bowel disease (IBD) refers to a group of diseases characterized by multifactorial pathology involving i) profound intestinal inflammation, ii) shifts in microbiome composition, and iii) intestinal tissue damage resulting in loss of gastrointestinal (GI) barrier integrity. The major subtypes of IBD are ulcerative colitis (UC) and Crohn’s disease (CD), with over 6 million people afflicted worldwide ^1,2^ resulting in impaired quality of life ^3–6^ and increased risk of cancer ^7^ and need for surgery ^8^. All current IBD therapeutics target inflammation; most are not specific to GI inflammation (e.g., anti-TNFα antibodies or JAK inhibitors), others are specific to GI inflammation (anti-a4b7 antibodies) ^9^. Regardless of their specificity, these treatments are all characterized by a so-called “therapeutic ceiling” in which only 10-20% of patients treated with current IBD treatments will achieve and maintain disease remission 1 year after treatment ^10^.

Furthermore, specific safety concerns (infection risk), high costs ($40,000 – $150,000 per year), and inconvenience (injection or infusion) limit treatment adherence and access to care, further degrading patients’ quality of life. Thus, a paradigm shift in IBD treatment is needed, with increased attention to the multifactorial causes of IBD, particularly environmental (e.g., gut microbiome), genetic (and epigenetic), and immune factors ^11–13^, towards delivery of more effective, safe, affordable, and convenient treatment for patients with IBD ^10^.

Among other biotechnological applications, bacterial extracellular vesicles (BEVs) are cell-secreted nanovesicles that have emerged as a promising therapeutic modality for IBD ^14^. BEVs from Gram-negative bacterial species have been developed as vaccines (Bexsero) but are not suitable for IBD applications due to high pro-inflammatory lipopolysaccharide (LPS) content ^15^. On the other hand, BEVs from Gram-positive species of probiotic/commensal strains found in the human microbiome lack LPS and are effective in preclinical models of IBD ^16^, as well as other inflammatory diseases. Often, these BEVs are derived from species of Gram-positive lactic acid bacteria (LAB), such as *Lactobacillus* spp. and *Bifidobacterium* spp.. Unlike current IBD therapies, these Gram-positive LAB BEVs can target novel therapeutic mechanisms to promote intestinal barrier integrity and immunomodulation, while retaining normal immune function, potentially owing to their evolved role as mediators of normal gut microbiome-host signaling. Thus, via these mechanisms, Gram-positive LAB BEVs have the potential to overcome therapeutic ceilings and drive more complete mucosal healing, which can promote durable remission of disease and mitigate long-term risks of cancer and need for surgery.

Despite the promise of Gram-positive LAB BEVs to improve IBD treatment, to date, their development has been limited due to i) low potency and ii) prohibitively low production yields caused by low biogenesis rates ^17^. These limitations are related - low potency necessitates high doses to achieve therapeutic effects, and high dose requirements exacerbate low production yields. Rigorous downstream purification processes can increase potency ^18^ but also result in product loss, whereas more crude methods contribute to batch variability during production, creating a major potential regulatory hurdle. Thus, engineered solutions to improve low yields and increase potency would greatly enable further development and translation of LAB BEV therapeutics for IBD. Toward this end, several approaches to increase BEV yields have been investigated. Environmental modifications or drug treatments have produced only modest 2-to 5-fold increases in BEV yields and could introduce regulatory concerns depending on the drug treatment used ^19–21^. Genetic deletion of membrane integrity proteins in Gram-negative bacteria such as *E. coli* can generate orders of magnitude increases in BEV yields, but at the expense of cell viability and BEV bioactivity ^22,23^, and homologous membrane integrity proteins do not exist in Gram-positive species of probiotics. Finally, extraction of BEVs from cell pellets using detergents or sonication is effective in Gram-negative bacteria ^24^, but Gram-positive bacteria are highly resistant to such approaches due to their distinct membrane architecture ^25^. Thus, while large BEV yield increases are possible in general, there is currently no robust system or method for generating meaningful increases in yields of LAB BEVs that might be most effective for IBD treatment.

In this study, we sought to develop such a system by focusing on four LAB species that have shown promising clinical efficacy as live cell formulations. In particular, an 8-strain combination of LAB (VSL#3) has established clinical efficacy in certain subtypes of IBD, i) mild-moderate ulcerative colitis, and ii) UC patients after ileal pouch-anal anastomosis surgery ^26–29^. We selected two strains from VSL#3 with clinical efficacy as single strain formulations in various GI diseases and symptoms, *Lactiplantibacillus plantarum* ^30–33^ and *Lacticaseibacillus paracasei* ^34^, and two other LAB strains also clinically effective, *Limosilactobacillus reuteri* ^35^ and *Lacticaseibacillus rhamnosus* ^36^. We compared BEV characteristics, in vitro anti-inflammatory activity, and efficacy in an acute DSS mouse model of IBD to identify a suitable cell source for genetic engineering. We then established proof-of-concept that LAB species, specifically *L. plantarum*, can be engineered to generate high amounts of BEVs via a genetically programmed hypervesiculation mechanism. We further showed that these high-yield BEVs retain their effectiveness in the IBD mouse model. Accordingly, the data supports continued preclinical development of hypervesiculating *L. plantarum* BEVs towards the translation of an effective, non-immunosuppressant, cost-effective, and convenient treatment for IBD that overcomes the therapeutic ceiling of conventional systemic immunotherapies.

## Methods

### Cell culture

*E. coli* Nissle 1917 (EcN was obtained from Mutaflor (Canada) and DH5a was obtained from New England Biolabs (Boston, MA). All lactic acid bacteria (LAB) strains were obtained through ATCC (*Lacticaseibacillus rhamnosus* GG (ATCC 53103), *Lacticaseibacillus paracasei* (ATCC 334), *Limosilactobacillus reuteri* F 275 (ATCC 23272), *Lactiplantibacillus plantarum* WCFS1 (ATCC BAA-793).

EcN and DH5a bacterial strains were cultured in LB media at 37°C at 250 rpm shaking for all experiments. All LAB strains were cultured at 37°C without shaking (static); for in vitro assays, LAB were cultured in De Man–Rogosa–Sharpe (MRS) media, and for bioactivity experiments, LAB were cultured in complete defined *Lactobacillus* media (LDMIIG) as in prior studies ^37^ except with two modifications: 1 g/l of Tween 80 was included and tryptophan increased to 0.2 g/L.

### BEV production

An overnight culture was started from a single colony. The overnight culture was then used to inoculate fresh media to commence BEV production. For *E. coli*, 200 ml of LB was inoculated at 1:150 and cultured for 16 h at 37°C and 250 rpm. For LAB, 400 ml of media (MRS or CDM) was inoculated at 1:50 and cultured for 24 h at 37°C without shaking. Then, the culture was depleted of cells and large debris by centrifugation at 10,000 g x 10 min (4°C), followed by a second centrifugation of the supernatant at 10,000 g x 20 min (4°C). Then, supernatant was filtered with a 0.45-um bottle top filter (Nalgene Rapid Flow PES; Thermo #169-0045) to generate clarified supernatant.

### Tangential flow filtration (TFF)

TFF was performed using a KrosFlo KR2i TFF system (Spectrum Labs, Los Angeles, CA, USA) equipped with a 300-kDa MWCO hollow fiber filter composed of a modified PES membrane (D02-E100-05-N; Spectrum Labs). Each filter was used no more than 5 times or discarded when permeate flow rate decreased, cleaned after each use by circulating 100 ml 0.5M NaOH for 30 min, and stored in 20% ethanol. Flow rate was set at 106 mL/min to maintain a shear rate of 4000 s^−1^ and backflow pressure was automatically adjusted to maintain a transmembrane pressure of 5 psi. Clarified supernatant was initially concentrated to 25 mL followed by diafiltration and then brought to a final concentration to ∼10 mL. Diafiltration volumes were scaled to 1.5 times the volume of clarified supernatant (600 ml for LAB and 300 ml for *E. coli*). Following TFF, samples were concentrated further to between 1-5 ml using 100 kDa ultrafiltration columns to achieve a final concentration of approximately 5E11 particles/ml. Finally, samples were sterile filtered with 0.2 uM PES syringe filters and stored at -20°C for no longer than 4 weeks before use.

### BEV characterization

Particle size and concentration were determined using NanoSight LM10 (Malvern Instruments) with Nanoparticle Tracking Analysis (NTA) software, version 2.3. Samples were diluted within the working concentration range of the NTA (particles/frame = 20-200) using PBS, and three 15-s videos were captured with a camera level set at 14. The detection threshold was set at 3 for all samples and replicates. Total protein concentration in BEV samples was determined using Bicinchoninic acid assay (BCA; 786-571; G-Biosciences). BEV proteome was analyzed by 12% SDS-PAGE followed by staining with SYPRO Ruby Protein Gel Stain (Thermo # S12001).

BEV morphology was assessed using negative stain TEM. First, 20 μL of BEV sample was mixed 1:1 with 4% EM-grade paraformaldehyde (157-4-100; Electron Microscopy Sciences) for 30 min at room temperature. Throughout the following steps, ultra-thin carbon-coated copper grids were placed, carbon film side down, on droplets of reagents positioned on a sheet of parafilm, with excess liquid blotted between each step. First, fixed BEVs were adhered to grids by floating grids on droplet of PFA-BEV mixture for 20 min. The BEV-adhered grid was then washed with PBS and floated on a droplet of 1% glutaraldehyde (in ×1 PBS) for 5 min. Next, the grid was washed five times with dH20 (2-min incubation per wash), and then negative stained using uranyl-acetate replacement stain (22405; Electron Microscopy Sciences) for 10 min. The grids were allowed to dry overnight before imaging on a JEOL JEM 2100 LaB6 TEM with a 200 kV accelerating voltage.

### Macrophage inflammatory assay (RAW264.7, dTHP1, CCK8)

RAW264.7 mouse macrophages were seeded at a density of 75,000 cells per well in a 48-well plate with DMEM containing 10% FBS and 1% penicillin/streptomycin. After 24 h, the cells were pretreated with media supplemented with one of the following: (i) ×1 PBS (six wells total), (ii) BEV-depleted conditioned media (permeate from TFF), (iii) 1 μg/mL dexamethasone as a positive control (Dex; D4902-25 MG; Sigma-Aldrich), or (iv) BEV groups isolated through different methods (TFF only, TFF + SEC, or TFF + HPAEC). All treatments were performed in triplicate, and BEV doses were normalized to particle content. After 24 h of pretreatment, the media was removed and 10 ng/mL LPS (resuspended in ×1 PBS; L4391-1MG; Sigma-Aldrich) was added to all wells, except for three PBS-pretreated wells (media-only group), to induce an inflammatory response. Conditioned media was collected 4 h after LPS stimulation, stored at - 80°C, and TNF-α levels were measured by ELISA within three days (DY410; R&D Systems). Additionally, cell lysate was collected in QIAzol and stored at -80C until RNA isolation within three days. For some experiments, prior to media and RNA collection, CCK8 viability assay was performed.

For human dTHP-1, a similar process was followed as RAW264.7 with a few modifications. First, THP-1 monocytes were differentiated into dTHP-1 macrophages by 48 h incubation with 20 nM phorbol 12-myristate 13-acetate (PMA). Then, cells were pretreated with BEVs as above, followed by stimulation with 500 ng/ml LPS and 20 ng/ml IFN-γ. Finally, 24 h after stimulation, conditioned media and/or cell lysate was collected for analysis of cytokine expression by ELISA.

### Murine acute DSS colitis model

All animal care and research were carried out using protocols approved and overseen by the University of Maryland and the University of Maryland IACUC (protocol R-MAR-23-11) in compliance with local, state, and federal guidelines. Male, 8–12-week-old, C57BL6J mice were purchased from The Jackson Laboratory. Colitis was induced by the addition of 2.5% (w/v) dextran sulfate sodium (DSS; MW ∼40,000; Chem-Impex) to autoclaved drinking water. Mice were weighed daily, and treatments were administered by oral gavage in a 200-ul volume. BEV doses were 2.5E9 particles/mouse/day, and live cell doses were 2.5E9 CFU/mouse/day. Sham treat control mice were administered vehicle (PBS). After colitis induction, DSS water was replaced with normal water constituting a “washout period”. At the end of study, colons were collected, their length measured from the ileocecal junction to rectum, and then prepared for histology by the Swiss roll method as previously described ^38^; mesenteric lymph nodes were collected and cells analyzed by flow cytometry; for some studies, RNA was isolated from the most proximal 1 cm of colon.

For some experiments, disease activity index was scored at the peak of disease on Day 4 and Day 5 according to the criteria in Table 1, as previously described^39^.

**Table 1.** Disease activity index scoring criteria.

|  | **Finding** | **Score** |
| --- | --- | --- |
| Stool consistency | Normal | 0 |
|  | Loose | 2 |
|  | Diarrhea | 4 |
| Blood in stool | None | 0 |
|  | Trace (hemocult positive) | 2 |
|  | Grossly positive | 4 |
| Weight loss | <1% | 0 |
|  | 1-5% | 1 |
|  | 6-10% | 2 |
|  | 11-15% | 3 |
|  | >15% | 4 |

### Flow cytometry

Harvested mesenteric lymph nodes (mLNs) were immediately placed on ice after collection. Then, mLNs were mechanically dissociated and passed through a 70-um cell strainer. Cells were washed with 1x PBS + 2% FBS (FACS buffer), then blocked with anti-CD16/CD32 antibody (TruStain FcX; clone 93; Biolegend #101320) for 15 min at 4°C. Following blocking, surface and LIVE/DEAD markers were stained for 30 min at 4°C. (Zombie Near IR Fixable Viability kit; Biolegend #423106). Then, cells were washed 2x with FACS buffer, and analyzed by flow cytometry within 12 h, or subjected to subsequent intracellular staining. For intracellular staining, cells were fixed and permeabilized using the Foxp3/Transcription factor staining buffer kit (eBioscience) and intracellular markers stained for 1 h at 4°C. Finally, cells were washed 2x and then analyzed by flow cytometry on a FACSCelesta (BD) with data analyzed using Flowjo software (Version 9, Treestar).

### RNA isolation and RT-qPCR analysis

RNA was extracted from either mouse proximal colon tissue or RAW264.7 cell lysate. The tissue was mechanically homogenized, and the cell lysate was collected using QIAzol. Total RNA was isolated with the RNeasy mini kit (QIAGEN, 74 104, Hilden, Germany) following the manufacturer’s protocol. Complementary DNA (cDNA) was synthesized from the total RNA using M-MuLV reverse transcriptase (New England Biosciences, M0253L, Ipswich, MA, USA) as per the manufacturer’s instructions. After cDNA synthesis, quantitative polymerase chain reaction (qPCR) was conducted using a QuantStudio 7 Flex qPCR system (ThermoFisher Scientific, 4 485 701, Waltham, MA, USA) and PowerTrack SYBR Green Master Mix (Thermo Scientific, A46109, Waltham, MA, USA). The primer sequences used for qPCR are provided in Table S1 (Supporting Information). mRNA transcript levels were quantified by the comparative Ct method, normalized to GAPDH expression (except in qPCR arrays), and expressed as fold changes in mRNA levels. For qPCR arrays, normalization was done using the average of four housekeeping genes (*gapdh*, *actb*, 18S rRNA, *eef1a1*). For mouse studies, RNA was isolated from individual mouse colons and converted to cDNA; then, equal masses of RNA were pooled from each mouse within a group to produce a single pooled cDNA sample for each group, RT-qPCR was performed on the pooled sample with three technical replicates.

### Histologic analysis

Paraffin embedded Swiss rolled colons were prepared as previously described and sectioned into 5-um thick slices and H&E stained. Stained sections of tissue were scanned into Aperio ImageScope and analyzed using methods described by Garcia-Hernandez et al ^40^. Briefly, lengths of normal, inflamed, and ulcerated tissue were measured, generating a Histological Colitis Score for each sample. Ulceration was defined as complete loss of the mucosa with disruption of colonic crypt architecture, and inflammation was defined as goblet cell density loss, crypt density loss, crypt hyperplasia, and/or presence of inflammatory infiltrate or edema^41,42^.

### Statistical analysis

All statistical analyses were performed using GraphPad Prism v9.0. Data from mouse studies were assessed for normality using the Shapiro-Wilk test. Tests used for statistical significance are indicated in figure captions. Statistical significance was defined as p < 0.05. All data are presented as mean ± standard deviation (SD), unless otherwise specified. Sample sizes were chosen based on prior studies. No data were excluded.

## Results

### LAB BEV characterization and assessment of media effects on production

We first characterized BEVs from the selected LAB species to assess their production and physical characteristics. LAB are typically cultivated in MRS media, a rich media-containing animal meat extract. Digested animal tissue can contain EVs, proteins, and other large debris. Some of these products can be co-isolated with BEVs and confound BEV proteomics and bioactivity analysis, not to mention their unsuitability for therapeutic biomanufacturing ^43,44^. Thus, we cultivated all strains of LAB in complete defined media (CDM) that does not contain animal products, and compared BEV characteristics to standard MRS. The size distribution of particles from all species was within the expected 20-200 nm range for BEVs (Figure 1A), and LAB cultured in MRS or CDM produced particles of similar size. The mode particle diameter was between 60-100 nm for all species except EcN, a probiotic *E*. *coli* species used as a control, which produced significantly larger particles with a mode size of 145 nm (Figure 1B). Of note, the other *E. coli* strain, DH5a, also produced a minor population of larger sized particles, which was insufficient to impact mode size. These findings are consistent with prior reports showing at least some Gram-positive species produce smaller diameter particles than Gram-negative *E. coli*^17^. These smaller particles with Gram-positive bacteria may reflect a biogenesis pathway that requires transit through the peptidoglycan cell wall. BEV yields were greater than 1E9 particles/ml culture for all species with up to 10-fold variability in yields between species; EcN produced the highest BEV yields at ∼2E10 particles/ml culture (Figure 1C). Surprisingly, despite normal growth in CDM, *L. plantarum* produced nearly 10-fold lower particle yields in CDM vs MRS.

**Figure 1.**
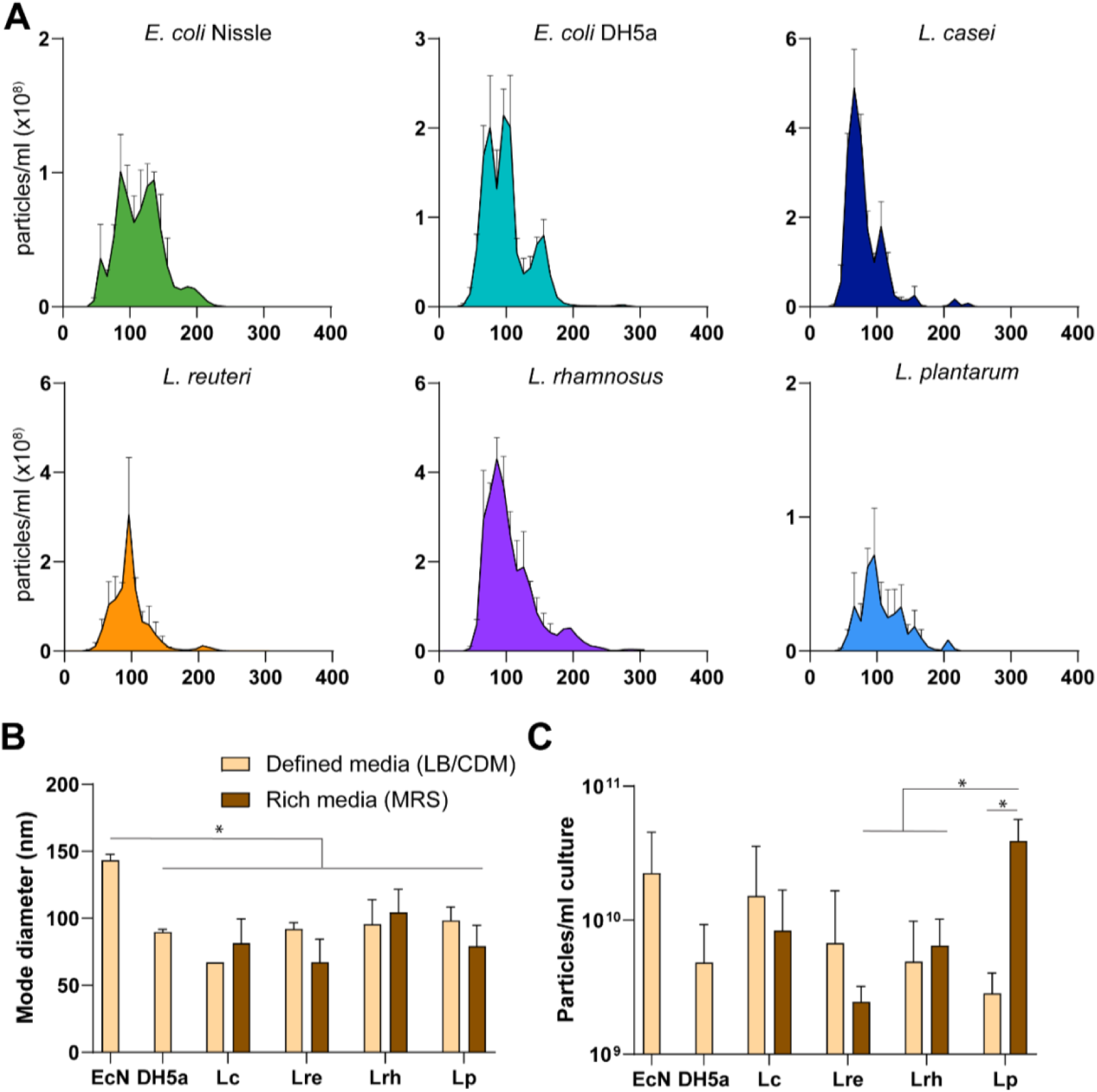
BEV size and production. **A)** Particle diameter distribution of BEV samples derived from different species of bacteria determined by nanoparticle tracking analysis (NTA) (error bars ± SEM), **B)** Mode particle diameter (nm ± SD) of BEV samples from different species of bacteria cultured in either defined media (LB or CDM) or rich media (MRS) determined by NTA, **C)** Total BEV yields (particles per ml culture ± SD) from flask culture determined by nanoparticle tracking analysis and normalized to volume of bacterial culture. Statistical analysis by one-way ANOVA with Tukey post hoc test * p<0.05.

### LAB BEVs reduce inflammatory responses from mouse and human macrophages

Next, we aimed to apply in vitro macrophage cytokine release assays to i) confirm BEV anti-inflammatory bioactivity, and ii) generate preliminary comparisons of anti-inflammatory potency between different species. Here, an ideal cell source would produce BEVs with the greatest anti-inflammatory potency, without signs of inducing toxicity in treated cells.

We screened BEV anti-inflammatory bioactivity with well-characterized in vitro macrophage cytokine release assays using both human and mouse cell lines. Our rationale for selecting this assay is as follows: i) IBD pathology is mediated in part by macrophage inflammatory responses, which recruits effector T cells to the intestine ultimately leading to profound inflammation and tissue damage, and ii) prior studies show macrophage inflammatory responses are regulated by LAB BEVs, potentially mediating therapeutic efficacy ^45^, iii) the macrophage cytokine release assay was previously shown to predict in vivo efficacy for mesenchymal stem cell derived EVs^46^.

BEVs from all selected LAB and *E. coli* species limited LPS-stimulated secretion of inflammatory cytokine, TNF-α, in mouse (Figure 2B) and human macrophages (Figure 2D). However, *L. paracasei* clearly displayed lower potency in both human and mouse assays, with significant reduction in TNF-α responses only observed at the highest dose in human dTHP-1 macrophages (Figure 2D). Among the LAB species, *L. rhamnosus* showed potentially more potent TNF-α suppression across both murine (Figure 2B) and human (Figure 2D) macrophages, though differences were relatively modest between BEVs from *L. rhamnoses*, *L. plantarum* and *L. reuteri*. From these data, we conclude that *L. paracasei* BEVs show reduced in vitro anti-inflammatory potency in LPS-stimulated macrophages compared to BEVs from *L. rhamnosus*, *L. plantarum* and *L. reuteri*.

**Figure 2.**
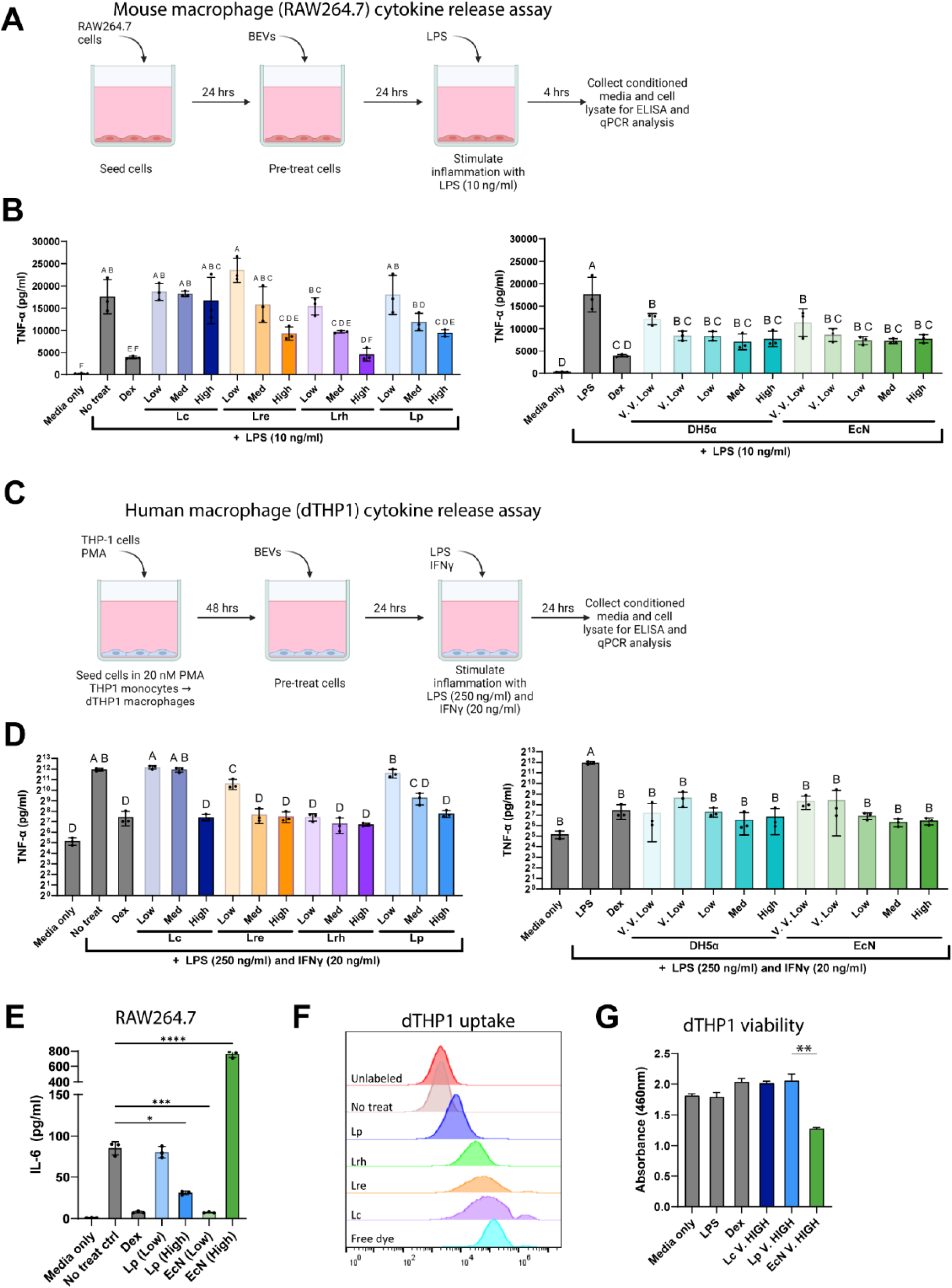
LAB BEVs exhibit anti-inflammatory effects in vitro. A-D) Mouse RAW264.7 (A-B) or human dTHP1 (C-D) macrophages pretreated with BEVs display reduced inflammatory TNF-α secretion following stimulation with 10 ng/ml LPS (for RAW264.7) or 250 ng/ml LPS + 20 ng/ml IFNγ (for dTHP1). Dosing (A-D): V.V. Low = 5E6 particles/ml, V. Low = 5E7 particles/ml, Low = 5E8 particles/ml, Med = 5E9 particles/ml, High = 5E10 particles/ml. **E)** RAW264.7 macrophages pretreated with BEVs from Gram-negative probiotic EcN BEVs or Gram-positive probiotic L. plantarum (Lp) BEVs followed by LPS stimulation; Dosing (E): Low = 1E9 particles/ml, High = 5E10 particles/ml. **F)** dTHP1 macrophages were treated with 5E9 particles/ml of BEVs that were previously covalently labeled with fluorescent Alexa Fluor 647;24 h later, cell uptake of fluorescently-labeled BEVs was analyzed by flow cytometry. Controls included BEVs not labeled with Alexa Fluor 647 (unlabeled), PBS/Vehicle treated cells (No treat), or free dye (Alexa Fluor 647 carboxylate 10 um) **G)** dTHP1 viability 24 h after treatment with Lc, Lp, or EcN BEVs (2e11 particles/ml) and LPS+IFNγ stimulation was assessed by CCK8 assay. **Abbreviations** in figures are as follows: Lp (L. plantarum), Lc (L. paracasei), Lre (L. reuteri), Lrh (L. rhamnosus), EcN (E. coli Nissle 1917), DH5a (E. coli DH5a), Dex (Dexamathosone). Statistical significance determined with one-way ANOVA and Tukey post hoc test. Groups marked with different letters are statistically significantly different, groups marked with the same letter are not significantly different. * p<0.05, ** p<0.01, ***p<0.001, ****p<0.0001. Error bars ± SD.

With respect to *E. coli*, BEVs from probiotic (EcN) and nonprobiotic (DH5a) strains reduced TNF-α responses at far lower (100-fold) doses than any LAB in both human and murine macrophages (Figures 2B and 2D, right panel). However, at higher doses, *E. coli* BEVs stimulated IL-6 release above levels of the no treatment control, whereas LAB BEVs reduced IL-6 at all doses tested (Figure 2E). Among many potential explanations for these divergent responses, Gram-negative *E. coli* BEVs contain LPS, a potent TLR4 agonist, whereas Gram-positive LAB BEVs lack LPS but do contain TLR2 agonists (e.g., lipoproteins). Differences in bioactivity may also be explained by differences in cell uptake; however, we observed all four LAB species were taken up by macrophages without correlation between uptake (efficiency or MFI) and reduction of TNF-α responses from macrophages (Figure 2F). Of note, BEVs are taken up by multiple pathways and we did not analyze specific uptake pathways ^47^. Finally, a key difference between LAB and *E. coli* BEVs was effects on cell viability at high doses (Figure 2G). Since LAB BEVs naturally lack LPS (endotoxin) which can cause toxicity, we hypothesized LAB BEVs would preserve cell viability at high doses. Indeed, LAB BEVs at high doses, up to 2e11 particles/ml (maximum feasible dose), did not reduce cell viability. On the other hand, *E. coli* (EcN) BEVs at high doses reduced cell viability.

### LAB BEVs show efficacy in acute DSS colitis murine model

For a more definitive comparison of therapeutic efficacy of BEVs from different LAB species, we evaluated candidate BEVs in a well characterized preclinical mouse model of IBD (ulcerative colitis) – acute dextran sulfate sodium (DSS) induced colitis ^48^. In the DSS colitis model, DSS ingested from drinking water forms nanoliposomal complexes in the intestines that are toxic to intestinal epithelial cells. Death of intestinal epithelial cells compromises barrier integrity, leading to increased flux of bacterial antigens from the microbiome into host tissues, triggering a massive inflammatory response. Ultimately, moderate to severe colitis develops which can be assessed by several endpoints including i) colitis-induced colon shortening, ii) disease activity index (DAI; weight loss, diarrhea, bleeding), iii) histology, and iv) cell populations and gene expression. For this screen, we focused on colitis-induced colon shortening owing to its widespread use to evaluate treatment efficacy in acute DSS murine models and its amenability to high throughput analysis. We supplemented this measurement by analyzing T cell populations in the mesenteric lymph nodes and colon tissue gene expression.

Following treatment, we found that BEVs from *L. plantarum* and *L. paracasei* significantly reduced colitis-induced colon shortening. Trends for reduced weight loss were also observed for these BEV treatment groups, but differences did not reach statistical significance (Figures 3A-B). Additionally, *L. plantarum* and *L. paracasei* BEVs reduced colitis-induced colon shortening significantly more than probiotic *E. coli* (EcN) BEVs, which had no significant effect on colon length at the 2.5E9 particles/mouse/day dose. Furthermore, *L. plantarum* and *L. paracasei* BEVs showed trends for superior efficacy that did not reach statistical significance compared to other LAB BEVs (*L. reuteri* and *L. rhamnosus*).

**Figure 3.**
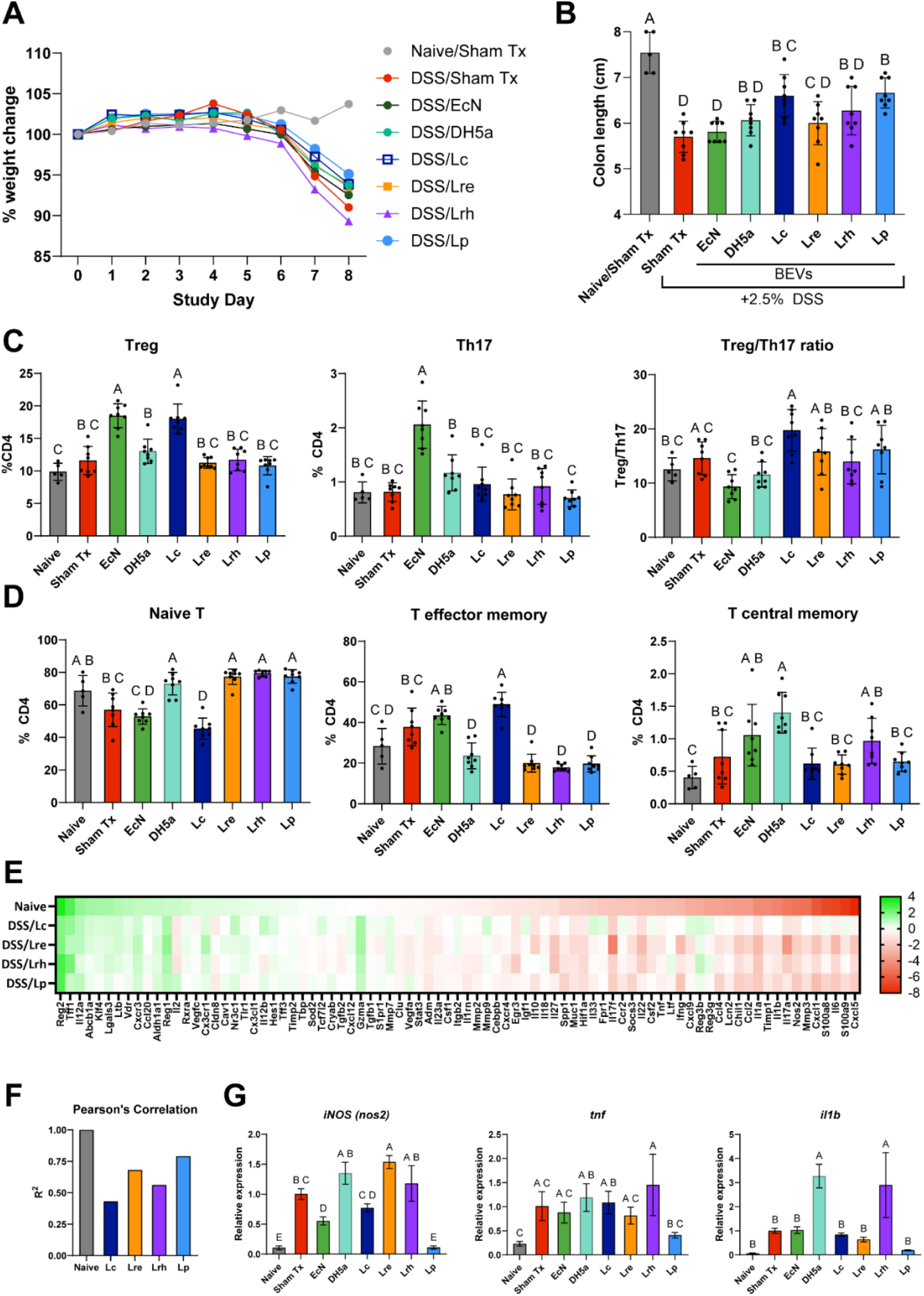
LAB BEVs reduce severity in acute DSS-induced colitis. Mice were treated with BEVs (2.5E9 particles/mouse/day) administered by oral gavage from Day 1-7; 2.5% DSS was administered in drinking water from Day 0 - Day 6, normal drinking water administered from Day 7-8 (washout period). **A)** Body weight changes of mice during acute DSS colitis relative to Day 0 weight, **B)** Colon length at study endpoint (Day 8), **C-D)** T cell populations and activation in mesenteric lymph nodes at study endpoint (Treg: CD4+CD8-CD25+Foxp3+, Th17: CD4+CD8-Foxp3-RORγt+; Naïve: CD4+ CD44-CD62L+; Effector: CD4+CD44+CD62L-; Central memory: CD4+CD44+CD62L+), E) Expression of genes associated with human ulcerative colitis in the proximal colon of mice was assessed by qPCR array and displayed as log2 fold change relative to sham (vehicle/PBS) treated mice with colitis. **F)** Pearsons correlation analysis was performed on colitis associated genes to assess correlations between treatment groups and naïve/healthy mice; R^2^ values closer to 1 indicated gene expression more closely associated with naïve/healthy mice. **G)** RT-qPCR of expression of specific genes involved in colitis in bulk colon RNA of mice. Abbreviations in figures are as follows: L. plantarum (Lp), L. paracasei (Lc), Lre (L. reuteri), Lrh (L. rhamnosus), EcN (E. coli Nissle 1917), DH5a (E. coli DH5a)). Statistical significance determined by one way ANOVA with Tukey post hoc test. Groups marked with different letters are statistically significantly different, groups marked with the same letter are not significantly different (p<0.05).

To explore potential mechanisms of the different LAB species and further validate conclusions of efficacy beyond colon length measurements, we evaluated T cell populations in mesenteric lymph nodes and gene expression in bulk colon tissue. We expected to find evidence of a regulatory T cell phenotype as previously reported during LAB BEV treatment in DSS colitis.

Instead, we found essentially unchanged Treg and Th17 populations for most LAB BEV treatments, with the exception of *L. paracasei* which modestly increased Tregs which may explain *L. paracasei* BEVs’s therapeutic efficacy (Figure 3C). We continued to explore potential mechanisms of *L. plantarum* BEV efficacy. Since no T cell polarization was appararent, we hypothesized that T cell activation was inhibited by *L. plantarum* BEV treatment. We evaluated antigen-dependent T cell activation in mLNs using the markers CD62L and CD44 to identify naïve, effector and central memory T cells. As expected, colitis increased T cell activation measured by reduced naïve and increased effector T cell populations (Figure 3D). With *L. plantarum* BEV treatment, we observed reduced T cell activation, as well as with *L. reuteri* and *L. rhamnosus* BEV treatment (Figure 3D). Consistent with *L. paracasei* BEVs induction of regulatory T cell reponses, we observed increased effector T cell activation with *L. paracasei* BEV treatment.

We found expected gene expression changes during induction of DSS colitis using a qPCR array of genes linked to human ulcerative colitis. DSS colitis was associated with increased pro-inflammatory genes (*il1b, tnf, ifng, nos2*), elevated surrogate markers of disease severity (*s100A8*, *s100A9*, and *lcn2*), and reduced expression of colitis-protective genes (*reg1*, *reg2*, *tff1*, *vdr*, and *abcb1a*) (Figure 3D). These gene expression changes defined a “colitis-associated gene expression profile”. Consistent with colon length data, global analysis of gene expression data using Pearson’s correlation analysis revealed *L. plantarum* BEVs best normalized the colitis-associated gene expression profile (R^2^=0.79, correlated to naive) (Figure 3F). Suprisingly, *L. reuteri* BEVs showed the 2^nd^ strongest correlation to naïve mice (R^2^= 0.68), despite no significant effect on colon length. Finally, *L. plantarum* BEV treatment reduced key inflammatoy markers to a greater extent than other BEV treatments (Figure 3G). Based on these outcomes, we identified *L. plantarum* as a strong candidate cell source for continued development of BEV IDB therapeutics.

### Validation and mechanistic analysis of L. plantarum BEV efficacy in acute DSS colitis model

To further validate the potential of *L. plantarum* BEVs for IBD therapy, we used the acute DSS mouse model as before, with the following treatment groups: i) two doses of *L. plantarum* BEVs (2.5E9 and 1E10 particles/mouse/day) and ii) a commercially available probiotic formulation previously shown clinically effective in randomized controlled trials (VSL#3), iii) single strain live *L. plantarum* cells, and iv) FDA-approved 1^st^ line drug for ulcerative colitis (5-ASA). These treatments were selected due to their predominant use in patients with mild-moderate ulcerative colitis – a likely initial application of therapeutic BEVs ^16^. The lower dose of BEVs (2.5E9 particles/mouse/day) was the same as prior studies, the higher dose (1E10 particles/mouse/day) was selected to assess efficacy and safety at supratherapeutic doses and guide determination of a therapeutic window.

As before, we found mice lost ∼5-10% body weight during colitis onset, without significant differences in weight loss across treatment (Figure 4A). Differences in colitis-induced colon shortening were significant, with all treatment groups except live *L. plantarum* cells showing significantly reduced colon shortening, indicative of reduced colonic tissue damage and therapeutic efficacy (Figure 4B). Interestingly, *L. plantarum* BEVs showed a trend for superior improvements in colon length compared to both live probiotic groups (Live *L. plantarum* cells and VSL#3) and 5-ASA, but the study was underpowered to detect significance. Finally, we included a co-treatment group of *L. plantarum* BEVs (2.5E9 particles/mouse/day) with 5-ASA (200 mg/kg/day) and found reductions in colitis-induced colon shortening with this combination, and trends for greater reductions compared to either treatment alone, suggestive of synergistic efficacy.

**Figure 4.**
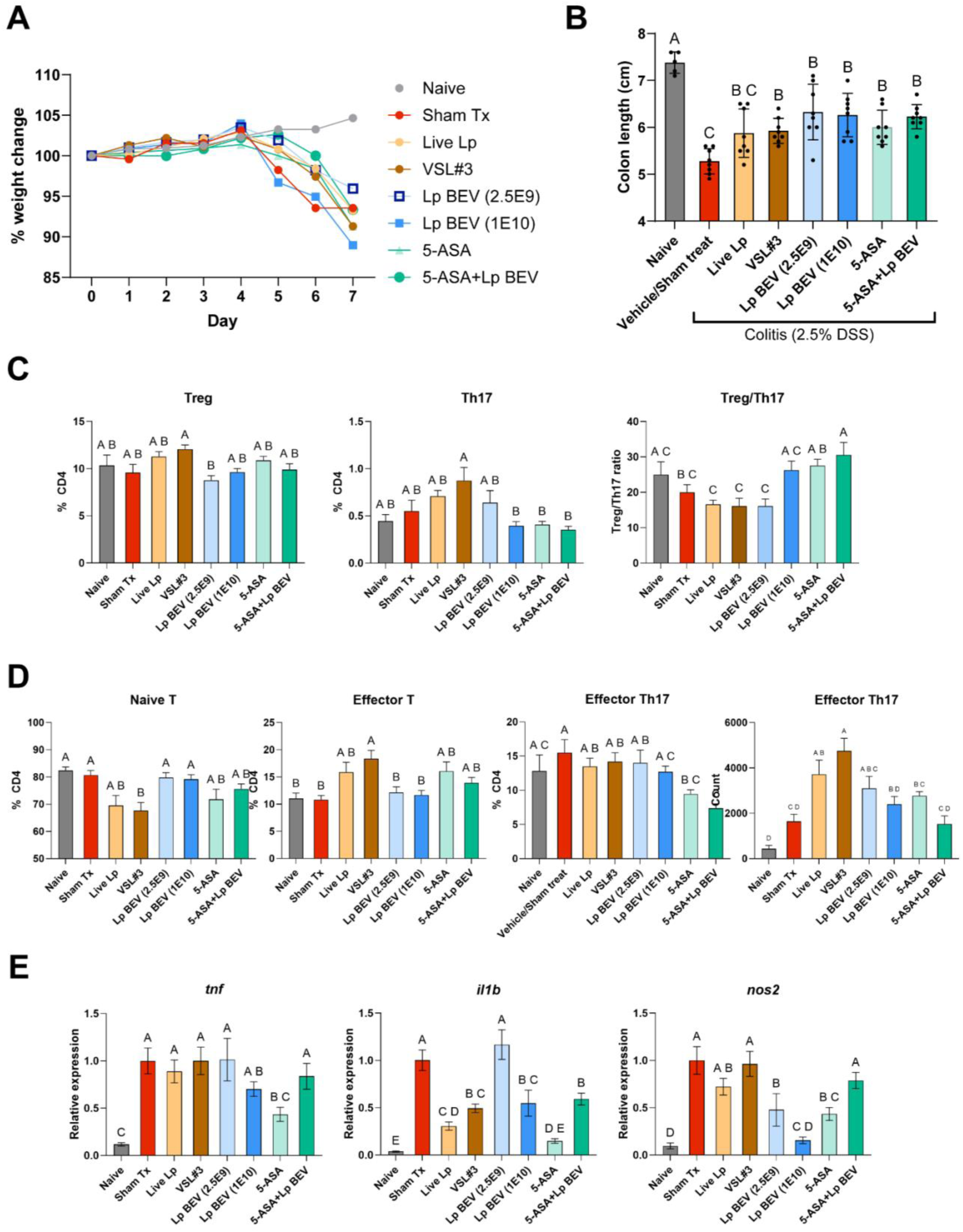
L. plantarum BEVs demonstrate similar efficacy compared to live probiotics and FDA-approved therapeutics in acute DSS-induced colitis, with potential distinct mechanisms. 2.5% DSS was administered in drinking water from Day 0-5, with normal H20 administered from Day 6-7. All treatments administered daily via oral gavage from Day 1-7; live *L. plantarum* and VSL#3 doses were 2.5E9 CFU/mouse/day, 5-ASA 200 mg/kg/day. For the *L. plantarum* BEV + 5-ASA group, the dose of BEVs was 2.5E9 particles/mouse/day. N=8 for each group except naïve where n=5. **A)** Weight change of mice as percent baseline weight, **B)** Colon length of mice at study endpoint (Day 7), **C-D)** Live single CD4+ T cell populations in mesenteric lymph nodes (±SEM). Treg: CD25+FOXP3+, Th17: CD25+FOXP3-RORγt+, Naïve T: CD44-CD62L+, Effector T: CD44+CD62L-, Effector Th17: CD44+CD62L-RORγt+. **E)** RT-qPCR analysis of gene expression in bulk colon tissue normalized to GAPDH expression; equal mass of cDNA from each mouse was pooled for RT-qPCR analysis, and 3 technical replicates performed. Statistical significance by one-way ANOVA with Tukey post hoc test, different letters above groups indicate statistical significance (p<0.05), the same letter above different groups indicate nonsignificant differences (p> 0.05).

Next, we evaluated T cell populations in mesenteric lymph nodes. Here, as in prior studies, *L. plantarum* BEVs did not polarize T cells towards a regulatory phenotype (Figure 4C). *L. plantarum* BEV treatment at 1E10 particles/mouse/day did significantly reduce populations of Th17 inflammatory cells. Likewise, 5-ASA and 5-ASA + *L. plantarum* BEV treatment reduced Th17 populations. Interestingly, both live probiotic treatments groups showed trends for increased T cell polarization, with a slight bias towards inflammatory Th17 subtype vs regulatory T cell (Treg) as depicted in Treg:Th17 ratio; we note differences were relatively modest (∼25-50% change) and did not reach statistical significance.

We again assessed the activation of T cells using CD44 and CD62L markers as in Figure 3. Here, consistent with live probiotics showing trends for increased Th17 polarization, we found significantly increased activated effector T cell populations following VSL#3 treatment, and a similar trend that did not reach statistical significance for live *L. plantarum* cells (Figure 4D). On the other hand, *L. plantarum* BEVs did not increase T cell activation, but unlike past studies, here we were unable to assess the inhibition of activation due to naïve and sham treated mice having similar levels of activated T cells, which may be explained by the short duration of the study or focus only on T cells in mesenteric lymph nodes.

Finally, we assessed expression of key inflammatory markers in the colon tissue of mice: i) *<u>tnf</u>*, an inflammatory cytokine target in IBD treatment, ii) *nos2*, an inflammation-induced producer of reactive nitrogen species that contribute to oxidative tissue damage, and iii) *il1b*, a potent inflammatory cytokine (Figure 4E). All these genes are primarily expressed by intestinal epithelial cells and myeloid immune cells and are implicated in IBD pathogenesis. We found *L. plantarum* BEVs reduced *nos2* expression significantly greater than both live probiotic treatments, and these effects were meaningful; live probiotics showed no reduction in *nos2* expression whereas *L. plantarum* BEVs showed dose-dependent reductions in *nos2* up to 90% below sham treated mice. This marker *nos2* bears clinical significance as a mediator of inflammation-induced tissue damage and warrants further investigation as a distinguishing mechanism between live probiotics and their secreted BEVs.

From these studies we concluded that: 1) *L. plantarum* BEVs show comparable preclinical efficacy in acute DSS murine models of IBD to clinically effective treatments for mild to moderate ulcerative colitis (VSL#3 and 5-ASA); 2) *L. plantarum* BEV treatment is potentially compatible in combination with current standard-of-care first-line therapy in mild to moderate ulcerative colitis (5-ASA); and 3) *L. plantarum* BEVs may have distinct and more favorable mechanisms of actions vs live probiotics based on reduced markers of oxidative tissue damage (*nos2*) and inflammatory Th17 cells.

### Generation of a hypervesiculating strain of L. plantarum for therapeutic BEV production

Upon confirmation that *L. plantarum* is a desirable cell source for production of therapeutic BEVs, we moved to solve the critical problem of low BEV production yields. Our approach was based on the premise that bacteria possess exceptional capacity for genetic engineering to augment or introduce desirable functions, and this can be applied for large scale increases in BEV yields. Thus, our goal was to genetically-program *L. plantarum* to mass produce BEV IBD therapeutics (“hypervesiculating” strain) and verify therapeutic efficacy of the resulting BEVs from the hypervesiculating strain.

Our design strategy was to harness a natural mechanism of BEV biogenesis: modulation of peptidoglycan crosslinking; this was achieved by transforming *L. plantarum* with a plasmid encoding a peptidoglycan remodeling enzyme under control of the inducible expression system, pSIP403 (Figure 5A) ^49^. We found this genetic circuit increased BEV production rates >66-fold, correlated with decreased cell density by OD600 measurements, with 22-fold increased total BEV yields in just 8 hours compared to conventional 24-hour culture (based on particle counts) (Figure 5B-C). An orthogonal method of BEV yield quantification (lipid quantification) showed even greater yield increases (Figure 5C). Further, not only were BEV yields dramatically increased, but also BEV purity dramatically increased based on an established BEV purity metric: particles per ug protein (Figure 5C) ^50,51^. These hypervesiculating *L. plantarum* BEVs displayed similar core proteome as normal BEVs, based on SDS-PAGE tracking of the most abundant protein bands in BEV samples (Figure 5D). Finally, hypervesiculating *L. plantarum* BEVs showed no significant difference in *in vitro* anti-inflammatory potency compared to normal BEVs in an LPS-stimulated mouse macrophage cytokine release assay (Figure 5E). From these data, we conclude that hypervesiculating *L. plantarum* BEVs show comparable structure, composition, and in vitro bioactivity to normal *L. plantarum* BEVs.

**Figure 5.**
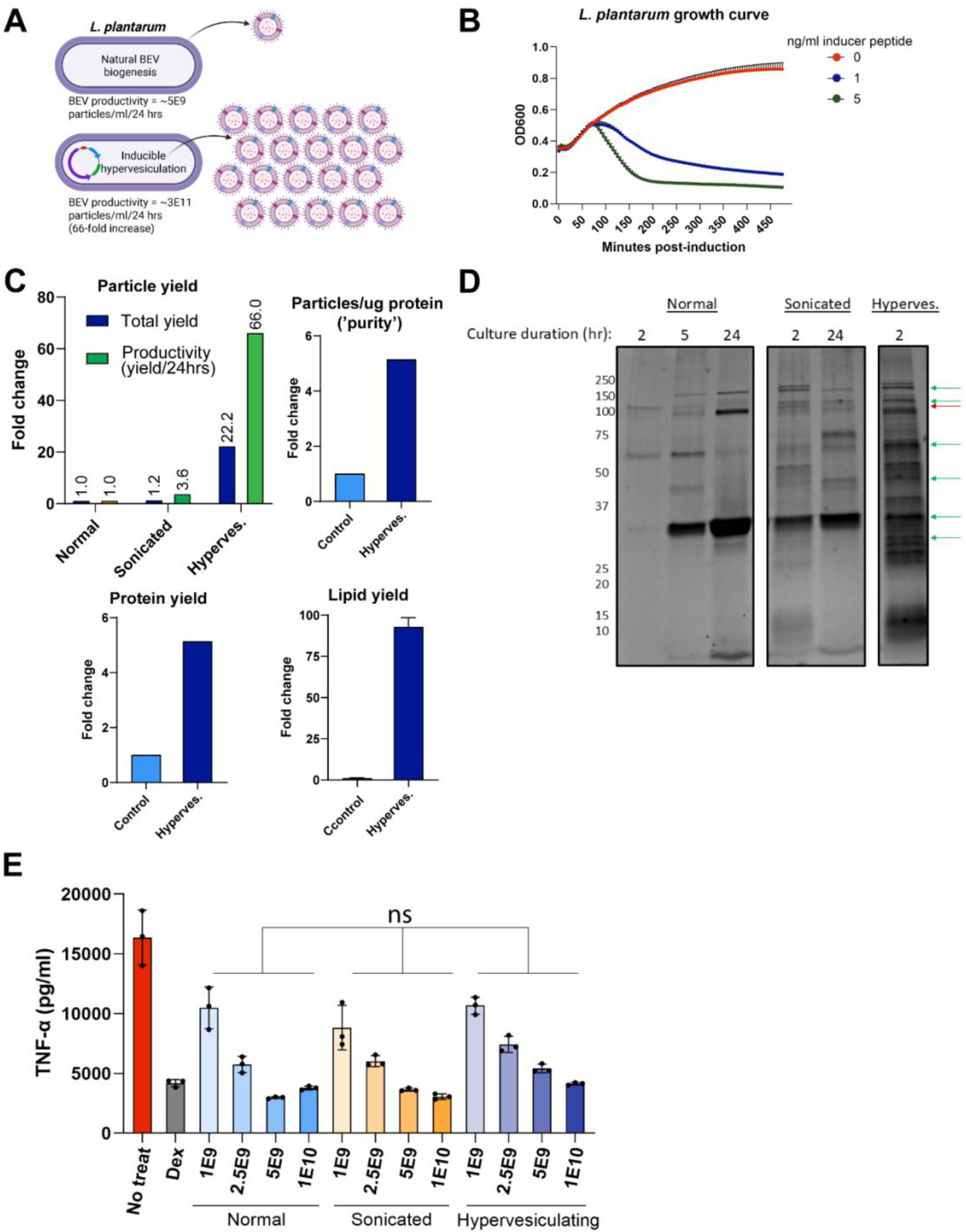
Engineering and characterization of hypervesiculating L. plantarum BEVs. A-C) Generation of a genetically-programmed hypervesiculating strain of L. plantarum with inducible activation (B) and increased BEV production yields (C), **B)** Hypervesiculating L. plantarum was cultured in 96-well plates at a starting OD ∼0.4 with inducer peptide (sppIP) added at time 0 at either 0, 1, 5 ng/ml and cell density monitored via OD_600_*, **C)** ∼2 h after addition of inducer peptide, BEVs were isolated from culture of either wild type L. plantarum (Ctrl) or hypervesiculating L. plantarum (Hyperves.) and yields determined by nanoparticle tracking analysis, total protein (bicinchoninic acid assay), total lipid (FM4-64 dye). Additionally, particle and protein yields were used to calculate a metric of BEV ‘purity’, particles/ug protein, and a sonicated control group was included which involved determination of BEV yields by NTA following sonication of L. plantarum cell pellet (derived from equal volume culture as other BEV groups) suspended in 30 ml of PBS (3 x 30 second pulses) and subsequent BEV isolation **D)** total protein analysis of BEVs by SDS-PAGE Coomassie stain reveals among the seven most abundant BEV proteins in normal BEVs, six were also detected in hypervesiculating L. plantarum BEVs (green arrows) and one was not detected in hypervesiculating L. plantarum BEVs (red arrow), E) RAW264.7 macrophages pretreated with BEVs followed by LPS stimulation do not show significant changes in secretion of the inflammatory cytokine TNF-α between normal BEVs, BEVs derived from sonication of cell pellet, or hypervesiculating L. plantarum BEVs, Statistical analysis by two-way ANOVA*.

*Hypervesiculating L. plantarum BEVs are effective in an acute DSS murine model of IBD* Finally, we sought to evaluate hypervesiculating *L. plantarum* BEV efficacy in the acute DSS colitis model, where normal *L. plantarum* BEVs had already shown efficacy (Figure 3). Mice were orally administered one of three treatments: i) hypervesiculating *L. plantarum* (live cells), ii) normal *L. plantarum* BEVs, iii) hypervesiculating *L. plantarum* BEVs; control mice were administered saline (sham treat). A reliable and accurate dose normalization method between cells and EVs is not possible, so we elected to use a dose of cells at the upper range of literature reports to allow preliminary comparisons of treatment efficacy and mechanisms^52–55^. Over the course of the study, all treated mice showed a trend for reduced weight loss compared to sham control, with ∼10% weight loss at Day 7 for sham vs. ∼5% for all three treatment groups (ns). We also found significantly reduced disease activity score (DAI) at the peak of disease (Day 5) for normal and hypervesiculating *L. plantarum* BEVs (Figure 6B). At the end of the study, we found significantly reduced colitis-induced colon shortening for all treatment groups (Figure 6C). There was no significant difference in colon length between the three treatment groups. Finally, blinded analysis of H+E-stained colon sections revealed reduced tissue ulceration in hypervesiculating *L. plantarum* BEV-treated mice vs sham (p=0.0599); whereas live *L. plantarum* cell-treated mice showed no improvement (Figure 6D). Hypervesiculating *L. plantarum* BEVs also showed a trend for greater reductions in tissue ulceration compared to normal *L. plantarum* BEVs.

**Figure 6.**
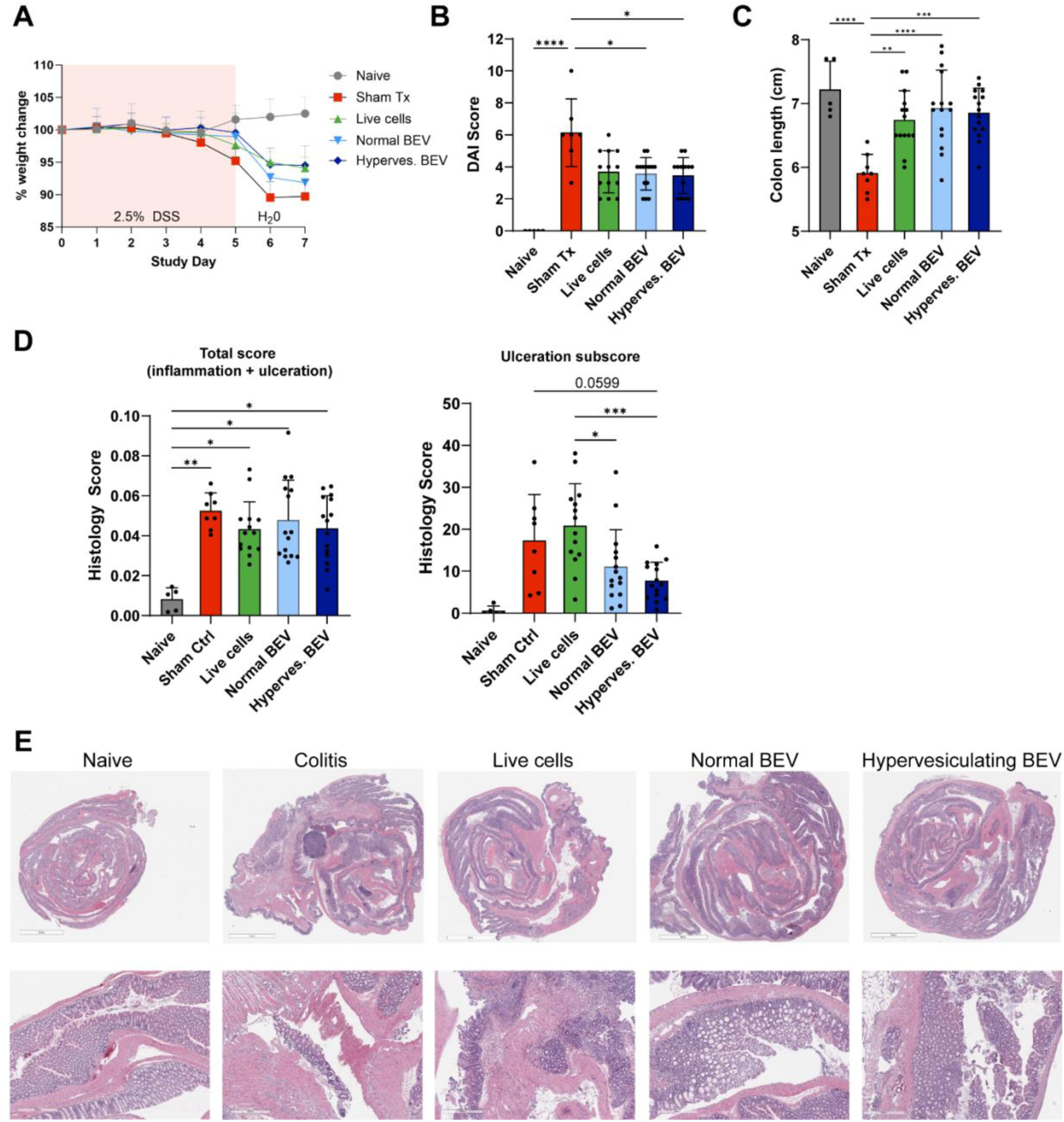
Hypervesiculating L. plantarum BEVs are effective in murine model of acute DSS-induced colitis. Mice undergoing acute DSS-induced colitis were treated with hypervesiculating L. plantarum BEVs (Hyperves. BEV), normal BEVs from L. plantarum, live L. plantarum cells, or vehicle (PBS; Sham Tx) daily from Day 1 – Day 6 via oral gavage, and 2.5% DSS was administered in drinking water from Day 0-Day 5. BEV doses were 2.5E9 particles/mouse/day, and the live cell dose was 2.5E9 CFU/mouse/day. N=15 for treatment groups, N=8 for Sham ctrl, N=4 for naïve. **A)** Body weight changes of mice during acute DSS colitis relative to Day 0 weight, **B)** Colon length at study endpoint (Day 7), **C)** Disease activity index (DAI) was assessed on Day 5. **D-E)** Histology analysis of H&E-stained colon section was performed by a blinded pathology team to assess tissue inflammation and tissue damage (ulceration) with representative images depicting hypervesiculating L. plantarum BEVs reducing tissue damage relative to control mice treated with saline and live cell treated mice. Statistical significance was determined by one- or two-way ANOVA and Tukey post hoc test. * p<0.05, ** p<0.01 ***p<0.001, ****p<0.0001.

## Discussion

There exist a myriad of potential bacterial cell sources to produce BEV IBD therapeutics. However, most are either poorly characterized, have unclear therapeutic relevance, or are not genetically tractable. Thus, our initial focus was to identify a suitable strain to implement engineering solutions to the most urgent hurdle facing BEV therapeutic’s clinical translation, low production yields. To this end, we took a combined rational and empirical approach involving first identifying genetically tractable cell sources that were applied in prior interventional human trials with at least encouraging results, followed by experimental evaluation of therapeutic activity of BEVs produced by these strains. Strengths of our approach include experimental design based on interventional human trial data and inclusion of both Gram-positive LAB species and Gram-negative *E. coli* species. Limitations of this approach include constraints on the number of cell source candidates imposed by the limited throughput of the acute DSS model. Additionally, further validation of hypervesiculating *L. plantarum* BEV efficacy in other preclinical models of IBD is required. Selection of candidate cell sources was complicated by most human trials utilizing probiotic formulations containing multiple species. To mitigate this, we also considered preclinical studies of BEVs produced by the candidate species. In general, species of Gram-positive LAB, such as *Lactobacillus* spp. and *Bifidobacterium* spp., met criteria for BEV cell source candidates. All four species of LAB examined here - *L. plantarum*, *L. paracasei*, *L. reuteri*, and *L. rhamnosus* - had previously shown efficacy as single strain live probiotics in human trials, and, for some, their BEVs showed efficacy in preclinical IBD models. An eight-strain cocktail of Gram-positive LAB denoted VSL#3, has shown consistent efficacy in certain human IBD subtypes (ulcerative colitis, and prevention of the post-UC surgery complication, pouchitis). VSL#3 contains *L. plantarum and L. paracasei*, in addition to other *Lactobacillus* and *Bifidobacterium* species. BEVs from *L. plantarum* and *L. paracasei* were previously shown to be effective in mouse models of IBD ^56^. Additionally, *L. reuteri* has shown promising clinical efficacy as a single strain formulation in preventing necrotizing enterocolitis in preterm infants ^57,58^, and also produces BEVs with anti-inflammatory effects in animals and human PBMCs ^59,60^. Finally, single strain *L. rhamnosus* was found to equally effective to standard of care treatment for ulcerative colitis ^61^, and also produces BEVs with anti-inflammatory effects in human PBMCs and efficacy in mouse models of IBD ^59,62^. Of note, for LAB species, we utilized the type strain of each species; strain level variation in BEV therapeutic activity could explain divergent results.

We also sought to draw comparisons of LAB BEVs to probiotic *E. coli* BEVs, that are under development as vaccines. This comparison is key, since technologies to increase BEV yields and load EcN BEVs with therapeutic protein cargo have already been developed. Live EcN has shown efficacy in human IBD equivalent to standard of care therapy ^63^, and EcN BEVs have efficacy in mouse models of IBD ^64^. Thus, *E. coli* BEV’s technology portfolio could support development of EcN BEV therapeutics for IBD.

To collect empirical data of BEV therapeutic activity, we employed a combination of in vitro and in vivo bioactivity screens. For the in vitro assay, we selected a well-characterized macrophage cytokine release assay. This assay was previously validated to predict in vivo efficacy of mesenchymal stem cell EVs in murine sepsis models, as macrophages are one of the main cell types involved in EV bioactivity. We found macrophages took up BEVs and BEV treatment resulted in dose-dependent reductions in secretion of the inflammatory cytokine, TNF-α, which is a therapeutic target in human IBD (UC and CD). Interestingly, we found EcN BEVs potently reduced TNF-α secretion with potency ∼100-fold greater than LAB BEVs. However, at higher doses EcN BEVs dramatically increased IL-6 secretion nearly seven-fold greater than purified LPS, whereas LAB BEVs suppressed IL-6 at all doses tested. Like TNF-α, IL-6 plays an important role in pathogenesis of chronic inflammatory and autoimmune diseases, and is a therapeutic target in rheumatoid arthritis, but not IBD. Thus, EcN induction of IL-6 responses from macrophages may detract from therapeutic efficacy in chronic inflammatory diseases with unclear relevance to IBD. Finally, at very high doses, EcN BEVs reduced cell viability, suggesting potential toxicity, likely due to its high LPS content. Several genetic strategies exist to detoxify LPS present on EcN BEVs, which could alter EcN BEV’s effects on IL-6 and in vitro cell toxicity.

We selected the acute DSS model to screen preclinical therapeutic efficacy of BEVs from candidate cell sources. We hypothesized that therapeutic BEVs would achieve therapeutic efficacy via innate immunomodulation and tissue repair. These processes are modeled in the DSS model where pathology is triggered by tissue damage (epithelial barrier) leading to innate immune activation. Other models, such as TNBS-induced colitis and T cell transfer, involve pathology mediated by the adaptive immune system (T cells). Key concerns of the DSS model are that certain effective IBD therapeutics are not effective in the model, and therapeutics are effective in this model but not in humans. Additionally, the gut microbiome is critical for DSS-triggered colitis. The gut microbiome of specific pathogen free laboratory mice housed in controlled conditions is poorly representative of the human microbiome’s potential role in dictating treatment responses. Future studies of BEVs in other models of IBD will be essential towards confirming the potential of BEV IBD therapeutics. In the acute DSS model, both *L. plantarum* and *L. paracasei* BEVs were effective at reducing colitis-induced colon shortening, whereas *L. plantarum* and *L. reuteri* BEVs most effectively normalized the colitis-associated gene expression profile. Thus, we concluded that *L. plantarum* BEVs are a suitable cell source to apply engineering to address low BEV production yields.

Prior to the study, we anticipated BEVs would show similar mechanisms of action as live probiotics. Interestingly, we found a stark contrast in efficacy and mechanisms with respect to markers of oxidative stress and histologic measures of mucosal tissue damage. Across several studies, *L. plantarum* BEVs dose-dependently reduced *nos2* (iNOS) expression by ∼90% whereas live *L. plantarum* cells, and a commercially available live probiotic formulation containing *L. plantarum* and 7 other Gram-positive LAB strains (VSL#3), showed no change despite high dosing. Additionally, we found live probiotics increased T cell activation and increased inflammatory T cell types (Th17), whereas *L. plantarum* BEVs either reduced or were neutral with respect to T cell activation and Th17 populations. Most importantly, *L. plantarum* BEVs reduced histologic measures of mucosal tissue damage (ulceration), whereas live *L. plantarum* cells showed no reduction, and even showed a slight trend for increased mucosal ulceration. These data suggest a divergent mechanism between *L. plantarum* cells and BEVs, potentially related to effects on oxidative tissue damage - critical pathology driving long term risks of cancer and surgery in IBD as well as reduced quality of life. While further validation is required in additional mouse models of IBD, this divergent mechanism reveals the potential of BEVs to show superior clinical efficacy versus live cell formulations.

Any BEV yield-generating method must generate orders of magnitude increased BEV yields to offer meaningful improvements in BEV biomanufacturing processes. Previous approaches to increase BEV yields in Gram-positive bacteria have included: i) cell disruption (e.g., French press) ^65^ or ii) environmental modification (e.g., pH, temperature, media agitation) ^19^.

Unfortunately, these approaches produce at best only 2-5-fold increased BEV yields and/or are not scalable. Low yields from cell disruption treatments can be explained by intrinsic resistance of Gram-positive bacteria to cell disruption, enabled by their thick peptidoglycan cell wall. Past experience with Gram-negative bacteria has shown genetic-based manipulation of BEV biogenesis, via decoupling of outer and inner membrane integrity by genetic knockout of membrane integrity proteins (e.g., tolA), can generate orders of magnitude increased BEV yields ^66–68^. These genetic technologies have already been applied in industrial biomanufacturing of Gram-negative BEVs for vaccine applications. However, homologous membrane integrity proteins do not exist in Gram-positive bacteria. Thus, our approach was to apply genetic control over Gram-positive BEV specific biogenesis mechanisms. Several mechanisms of BEV biogenesis in Gram-positive species have been proposed and reviewed ^69^: i) cytosolic turgor pressure ^70^, ii) outward membrane curvature, iii) decreased crosslinking of the peptidoglycan cell wall ^20,71^, and iv) membrane interactions with certain proteins, most notable the phenol soluble modulin peptides in *S. aureus* ^72^.

We found that manipulation of peptidoglycan cell crosslinking generated rapid and robust increased BEV yields, with 66-fold increased BEV production rate. A key question is whether hypervesiculating *L. plantarum* BEVs have preserved composition and bioactivity relative to normally-generated BEVs. Indeed, prior studies revealed hypervesiculating strains of Gram-negative *E. coli* (tolR knockout) produce BEVs with altered morphology, composition, and reduced capacity for epithelial cell uptake ^73^. These changes may compromise their therapeutic efficacy. We found encouraging evidence that our genetically-programmed hypervesiculating *L. plantarum* strain does not suffer from such limitations. First, therapeutic efficacy in the acute DSS model is retained, with nonsignificant differences in colitis-induced colon shortening, weight loss, DAI, and histologic score between normal and hypervesiculating *L. plantarum* BEVs. Second, protein composition analysis by SDS-PAGE reveals that the most abundant proteins in normal *L. plantarum* BEVs are also present in high levels in hypervesiculating *L. plantarum* BEVs. Further investigation of therapeutic efficacy in other mouse models (e.g., chronic DSS, IL-10 knockout, adoptive T cell transfer) could bolster these findings.

## Conclusions

Overall, the data generated here support the conclusion that hypervesiculating *L. plantarum* BEVs are effective in the acute DSS murine model of IBD, potentially via mechanisms involving enhancement of mucosal tissue repair/protection and immunomodulation, especially with respect to oxidative tissue damage. As such, hypervesiculating *L. plantarum* BEVs are a promising potential novel IBD therapeutic that solves a key biomanufacturing limitation of EV therapeutics related to low production yields.

## Acknowledgments

The authors acknowledge the following funding sources for support of the work reported: NIH HL141611 (to SMJ). This work was supported by Merit Review Award BX004890 from the United States (U.S.) Department of Veterans Affairs Biomedical Laboratory Research and Development Program (to JPR) and the National Institutes of Health, National Institute of Diabetes and Digestive and Kidney Diseases (award number T32 DK067872 to JPR). The contents do not represent the views of the U.S. Department of Veterans Affairs or the United States Government. This work was also supported in part by a Center of Excellence in Regulatory Science and Innovation grant to the University of Maryland from the US FDA 5U01FD005946 (to WEB). This article reflects the views of the authors and should not be construed to represent the US FDA’s views or policies.

